# Effect of Adolescent Stress on Adult Morphine-Induced Behavioral Sensitization is Dependent Upon Genetic Background

**DOI:** 10.1101/2021.01.20.427504

**Authors:** Helen M. Kamens, Carley N. Miller, Jasmine I. Caulfield, Dana Zeid, William J. Horton, Constanza Silva, Aswathy Sebastian, Istvan Albert, Thomas J. Gould, Diana Fishbein, P. Sue Grigson, Sonia A. Cavigelli

## Abstract

Deaths related to opioid use have skyrocketed in the United States, leading to a public health epidemic. Research has shown that both biological (genes) and environmental (stress) precursors are linked to opioid use. In particular, stress during adolescence – a critical period of frontal lobe development – influences the likelihood of abusing drugs. However, little is known about the biological mechanisms through which adolescent stress leads to long-term risk of opioid use, or whether genetic background moderates this response. Male and female C57BL/6J and BALB/cJ mice were exposed to chronic variable social stress (CVSS) or control conditions throughout adolescence and then tested for morphine locomotor sensitization or morphine consumption in adulthood. To examine possible mechanisms that underlie stress-induced changes in morphine behaviors, we assessed physiological changes in response to acute stress exposure and prefrontal cortex miRNA gene expression. Adolescent stress did not influence morphine sensitization or consumption in BALB/cJ animals, and there was limited evidence of stress effects in female C57BL/6J mice. In contrast, male C57BL/6J mice exposed to adolescent CVSS had blunted morphine sensitization compared to control animals; no differences were observed in the acute locomotor response to morphine administration or morphine consumption. Physiologically, C57BL/6J mice exposed to CVSS had an attenuated corticosterone recovery following an acute stressor and downregulation of twelve miRNA in the prefrontal cortex compared to control mice. The specificity of the effects for C57BL/6J versus BALB/cJ mice provides evidence of a gene by environmental interaction influencing opioid behaviors. Long-term differences in stress reactivity or miRNA expression suggests two possible biological mechanisms to evaluate in future research.

## 2 Introduction

The use of opioids is a widespread public health crisis. Between 1999 and 2018 opioid overdoses accounted for 440,000 deaths in the United States (Wilson et al., 2020). This number has been on the rise with over 46,000 Americans losing their lives to an opioid overdose in 2018 alone (Wilson et al., 2020). These deaths arise from the 10.3 million individuals who report misusing opioids each year (Substance Abuse and Mental Health Services Administration, 2019). Given the prevalence of opioid abuse in the United States, research into the mechanisms that increase the risk of opioid use is important for combating this public health crisis.

Opioid abuse is influenced by multiple factors. Human genetics studies have demonstrated that genes influence opioid use and dependence with heritability estimates ranging from 0.52 – 0.76 (van den Bree et al., 1998; Kendler et al., 1999, 2000; Sun et al., 2012). In this work, the use of different types of opioids (e.g. morphine, oxycodone, fentanyl) is not differentiated. However, research with inbred panels of mice have confirmed a genetic influence for individual opioids, including morphine, with heritability estimates of morphine behavioral responses ranging from 0.26 - 0.44 (Bergeson et al., 2001; Kest et al., 2002; Liang et al., 2006). Although these findings support genetic influences on opioid use and responses, environmental factors also impact opioid use.

One environmental influence linked to opioid use is exposure to stress. Research in animal models has demonstrated that exposure to social stressors can influence opioid behaviors (Neisewander et al., 2012). Limiting results to only mouse models provides a more consistent pattern. In male C57BL/6J mice exposed to chronic social stress in adulthood increased morphine preference in a 2-bottle choice experiment immediately after the stressor, but the effect was not long lasting (Cooper et al., 2017). Also, exposure to social defeat in adult male OF1 mice reinstated morphine conditioned place preference following extinction (Ribeiro Do Couto et al., 2006). These data suggest that adult social stress can influence morphine behaviors; however, there is limited data on how social stress during adolescence influences behavior related to drug use. To our knowledge there has been one study that examined the effect of an adolescent stressor, social isolation for 30 days, on the rewarding effects of morphine in mice. In male NMRI, male morphine conditioned place preference was eliminated in animals raised in isolation (Coudereau et al., 1997). What has not yet been fully characterized during this developmental period are the mechanistic links that can lead to opioid use and if these effects are also observed in female mice.

Adolescence is a critical time of brain and social development (Spear, 2013) and stress during this time generally leads to increased substance use in both humans and in animal models (Dube et al., 2003; Andersen and Teicher, 2009). The prefrontal cortex (PFC), which is responsible for executive functions, such as decision making, impulse control, and working memory has important connections with limbic structures involved in drug use. This brain region undergoes numerous structural and functional changes during adolescence. These dynamic changes open a window for environmental factors, particularly stress, to modulate processes associated with drug use (Jankord et al., 2011). In particular, the PFC is involved in regulating the hypothalamic-pituitary-adrenal (HPA) axis which matures during adolescence and has been implicated as one pathway by which social stress may affect opioid behavioral responses in adult rats (Deroche et al., 1994, 1995). One long-lasting molecular change found as a result of adolescent stress is miRNA expression patterns (Rao and Pak, 2016; Liu et al., 2017; Xu et al., 2017). These small RNA molecules regulate the expression of messenger RNA molecules, and thus could lead to wide scale changes in gene expression profiles, neuronal function, and altered drug behaviors.

The current study aims to investigate both environmental and genetic influences on opioid behaviors in a mouse model. We examined the influence of adolescent social stress on opioid behaviors in two different inbred mouse strains. The C57BL/6J and BALB/cJ strains were chosen because they are sensitive to long-term alterations in drug behaviors following chronic social stress in adolescence (Caruso et al., 2018c, 2018b, 2018a). In prior research, adult social stress (isolation or social defeat) led to increased voluntary morphine consumption (Alexander et al., 1978; Raz and Berger, 2010; Cooper et al., 2017). Thus, we hypothesized that chronic variable social stress (CVSS), a model of adolescent social stress that includes repeated cycles of social isolation and social re-organization, would increase morphine consumption and behavioral sensitization to morphine. To identify a potential mechanism by which adolescent stress alters morphine behaviors, we measured corticosterone (CORT) responses to an acute stressor and miRNA expression patterns in the PFC. As above, this brain region was chosen because it is known to be susceptible to stress manipulations and to influence drug behaviors (Andersen and Teicher, 2009). Additionally, prior work we have shown long-term alterations in PFC excitatory transmission following CVSS (Caruso et al., 2018a).

## 3 Materials and Methods

### 3.1 Animals

Male and female BALB/cJ and C57BL/6J mice (The Jackson Laboratory, Bar Harbor, ME, USA) were bred at The Pennsylvania State University. On postnatal day (PND) 21 offspring were weaned into groups of 2-3 same sex littermates housed in polycarbonate cages (28 cm × 17 cm × 12 cm) lined with corn-cob bedding. The animals were maintained on a 12-h light-dark schedule (lights on at 0700 h) in a temperature- and humidity-controlled facility with *ad libitum* food and water available. All procedures were approved by The Pennsylvania State University IACUC committee.

### 3.2 Drugs and Solutions

For morphine consumption, morphine sulfate pentahydrate was obtained from the NIDA Drug Supply Program (Bethesda, MD), and saccharin sodium salt and quinine hemisulfate salt were obtained from Sigma-Aldrich (St. Louis, MO). Morphine, saccharin, and quinine were diluted in tap water for the consumption study (see below). For locomotor sensitization, morphine sulfate pentahydrate (Sigma-Aldrich, St. Louis, MO) was diluted in physiological saline (0.9% NaCl; Baxter) and injected intraperitoneally in a 10 ml/kg volume. The two suppliers of morphine were based on differences in funding sources between projects.

### 3.3 Chronic Variable Social Stress (CVSS)

To minimize litter effects, littermates were evenly distributed between the CVSS and control groups at weaning. CVSS mice underwent the stress protocol during adolescence (PND 25-59) as previously described (Caruso et al., 2017, 2018c, 2018b, 2018a). Following weaning mice sat undisturbed with littermates (PND 21-25). On PND 25, CVSS mice were individually housed for 3 days followed by social rehousing with 1-2 unfamiliar cage mates for 4 days. This procedure was repeated for 5 consecutive weeks. On PND 59, CVSS mice were rehoused with their original cage mates from weaning until behavioral testing or sacrifice. For control mice, cage mates remained the same from weaning to the end of the study. To equate handling between the control and CVSS conditions, each time CVSS mice were transferred to a new cage for individual or social rehousing, control mice were also moved to new cages. Table 1 details the number of CVSS and control animals tested for each behavioral outcome described in detail below. Importantly, all behavioral and physiological outcomes were examined in independent groups of animals.

**Table 1.**
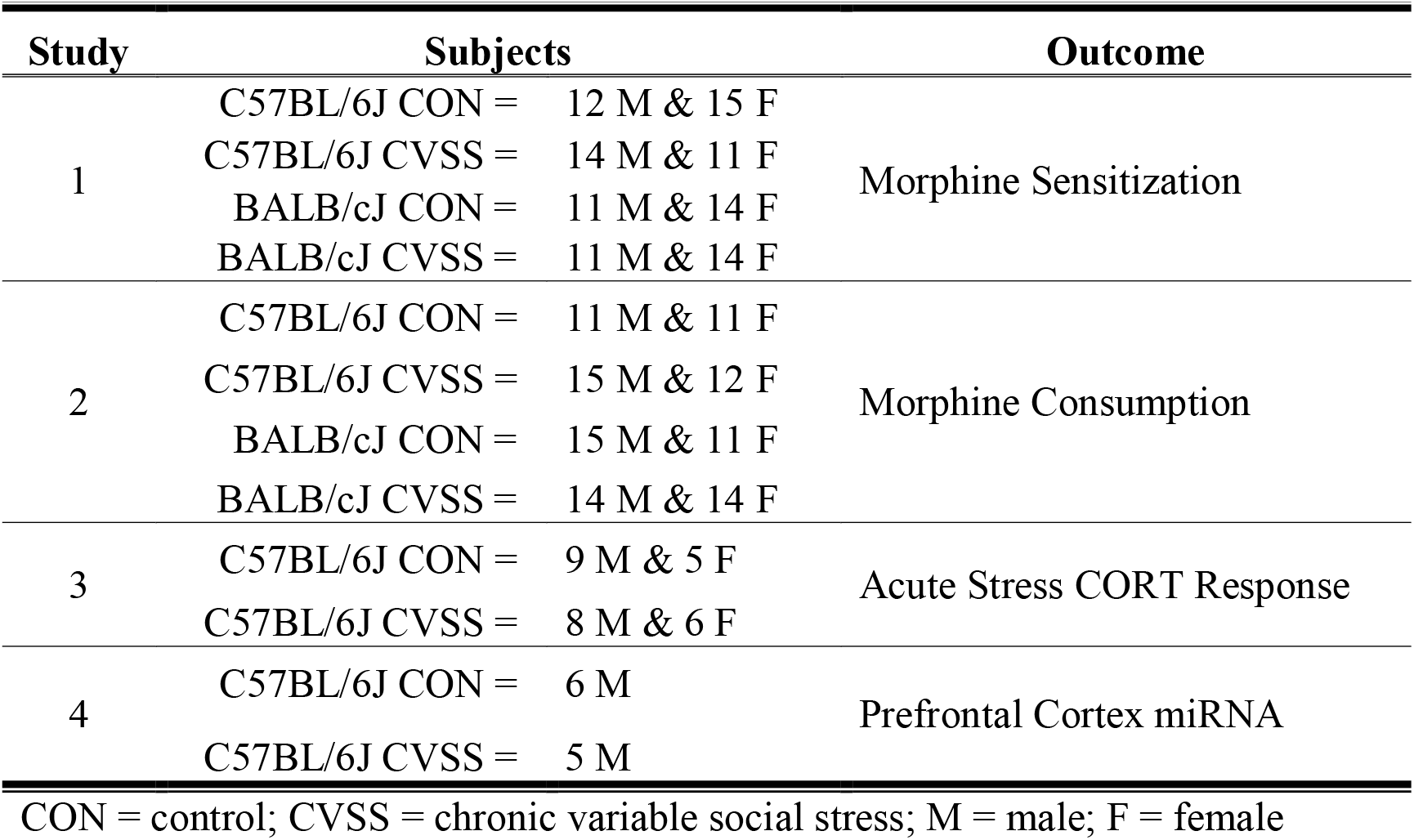
Overview of experimental design.

### 3.4 Methods of Behavioral and Physiological Outcomes

#### 3.4.1 Morphine Sensitization

Approximately 1 week after CVSS was complete, adult mice (PND 65-72) were tested for morphine-induced locomotor sensitization using published methods (Phillips et al., 1994b; Kamens et al., 2005). To achieve the desired sample size, two sequential cohorts were used that were counterbalanced for strain, sex, and stress condition. This age was chosen because our prior work demonstrated changes in drug behavior and physiological responses at this time (Caruso et al., 2017, 2018c, 2018b, 2018a). Testing was conducted between 0900-1300 h. On test days, all animals were moved to the experimental room and allowed to acclimate for at least 30 min. To acclimate mice to the test chambers, on days 1-2 all mice received an acute injection of saline immediately prior to being placed into a Superflex locomotor activity chambers (Omnitech Electronics, Inc., Columbus, OH). On day 3, animals were randomly assigned to the chronic saline or chronic drug group. Chronic saline mice received saline and chronic drug mice received 100 mg/kg morphine on days 3, 5, 7, and 9 immediately before being placed in the activity chamber. On day 11, mice in both the chronic saline and chronic drug conditions received 100 mg/kg morphine. This high dose of morphine was chosen because it has been shown to elicit the development of behavioral sensitization in adolescent and adult C57BL/6J animals (Koek, 2014) and was used in prior work on long term effects of adolescent stressors in male NMRI mice (Coudereau et al., 1997). On each day, locomotor activity was measured for 30 min and data were recorded as centimeters (cm) travelled. Three scores were calculated to examine acute morphine locomotor stimulation and sensitization within the chronic saline or chronic drug groups. Acute morphine stimulation was calculated as Day 3 – Day 2 locomotion in the chronic drug group and Day 11 – Day 2 locomotion in the chronic saline group. Sensitization in the chronic drug group was measured as Day 11 – Day 3 locomotion (Kamens et al., 2005). In each strain, 12-14 mice were tested in each of the four stress X drug groups (Control-Chronic Drug, Control-Chronic Saline, CVSS-Chronic Drug, CVSS-Chronic Saline), with approximately equal numbers of males and females in each group.

#### 3.4.2 Morphine Consumption

With a separate cohort of adult mice (starting PND 65-68) 2-bottle choice morphine consumption was tested using published methods (Eastwood and Phillips, 2014). Mice were individually housed and provided with two 25-ml graduate cylinders fitted with sipper tops. To acclimate mice to the new test environment, water was available for 2 days in both tubes. After acclimation, water tubes were replaced with one tube containing 0.2% saccharin plus morphine while the other tube containing 0.2% saccharin plus quinine. Quinine was provided as the alternative solution to account for the bitter taste of morphine following published protocols (Eastwood and Phillips, 2014). Drug solutions were available 24 hours a day. Morphine/quinine concentration increased every 4 days (morphine: 0.3, 0.7, 1 mg/ml; quinine: 0.2, 0.4, 0.55 mg/ml). Body weight was measured each time the drug concentration changed. The side of the cage where morphine was presented was alternated every other day to prevent the development of a side preference. Morphine consumption (mg/kg), quinine consumption (mg/kg), and total fluid consumption (ml) were calculated and used as dependent variables. For each drug concentration, the average of day 2 and 4 consumption values were used to assess stable drug intake (Phillips et al., 1994a). For each strain, 22-28 mice were tested from each stress condition, evenly split between males and females.

#### 3.4.3 Acute Stress

Our data indicated that C57BL/6J, but not BALB/cJ mice, showed different behavioral responses to morphine following adolescent stress (see results), therefore the remaining measures were limited to this strain. Adult C57BL/6J mice (PND 81-83) were used to measure the glucocorticoid response to an acute stressor. We used a standard restraint stress protocol administered between 0900-1100 h. Mice were removed from their home cage and placed in a broom-style restrainer. The tip of the tail (<1 mm) was removed and a baseline blood sample was collected within 3 min of initial disruption. Mice remained in the restrainer for 15 min then were placed into individual holding cages. At 30 and 90 min after initial placement in the restraint tube, mice were briefly returned to restrainers for an additional blood sample collection. These time points were chosen to capture peak (30 min) and recovery (90 min) corticosterone (CORT) levels following an acute stressor (Romeo, 2010). Stress reactivity was defined as 30 min CORT – baseline CORT and recovery was defined as 90 min CORT – 30 min CORT (Sapolsky et al., 1983). These two scores were used as dependent variables. Fourteen CVSS and 14 control animals were tested in this protocol including both sexes. Blood samples were collected into heparinized capillary tubes and immediately stored on ice. Plasma was obtained by centrifuging samples for 15 min at 10,000g at 4°C then stored at -80°C. CORT was measured using a commercial [^125^I] radioimmunoassay kit (MP Biomedicals, Solon, OH, USA) following manufacturer guidelines (Caruso et al., 2018b, 2018a). Intra- and inter-assay coefficients of variation were 7.8 and 0.8 (for low control) and 7.5 and 7.5 (for high control).

#### 3.4.4 Prefrontal Cortex (PFC) miRNA Gene Expression

To examine adolescent stress-induced alterations in miRNA gene expression 11 male C57BL/6J mice, unexposed to opioids, were used. Only C57BL/6J males were examined because they exhibited a change in morphine sensitization following CVSS (see results). Following CVSS, these mice remained undisturbed (i.e. no behavioral testing) until PND 70 when they were euthanized by cervical dislocation and brains removed. A mouse brain matrix was used to obtain 1 mm slices. The PFC was punched from the slice with a blunt needle that captured both hemispheres and immediately place into RNAlater. Tissue was homogenized and RNA was extracted using the RNeasy® Plus Universal Mini Kit (Qiagen, Valencia, CA, USA) following the manufacturers’ guidelines for the isolation of total RNA including small RNAs. RNA quality was determined using the RNA Integrity Number (RIN) with an Agilent 2100 Bioanalyzer (Santa Clara, CA, USA).

RNASeq libraries were prepared using the TruSeq Small RNA Library Construction Kit (Illumina, San Diego, CA, USA). The resulting libraries were quantified and pooled for sequencing on an Illumina HiSeq to obtain 50 bp reads. On average we obtained 7 million reads per sample. Library preparation and sequencing was conducted in the Penn State Genomics Core Facility, University Park, PA. Sequencing data are available from the NCBI GEO database (experimental series accession number: to become available upon publication).

Sequencing reads were mapped to the mouse genome (*Mus musculus* mm10) using ShortStack (Johnson et al., 2016). miRNA annotations were downloaded from miRBase and reads mapped to miRNAs were counted in ShortStack using the default settings. Differential expression between groups was performed using DESeq2 (Love et al., 2014). *P*-values were corrected for multiple comparisons using a Benjamini-Hochberg false discovery rate of 5% (Benjamini and Hochberg, 1995).

Differentially expressed miRNAs were analyzed for predicted mRNA targets and associated biological pathways using the DIANA toolkit (available at http://diana.imis.athena-innovation.gr/DianaTools/) and Ingenuity Pathway Analysis (IPA) software (content version 52912811; Krämer, Green, Pollard, & Tugendreich, 2014). The number of miRNA-gene relationships was predicted using the DIANA microT-CDS tool, v.5 (Paraskevopoulou et al., 2013), which computes miRNA-mRNA interaction scores for predicted targets in mRNA 3′-UTR and coding sequence (CDS) transcript regions. The microT-CDS score threshold was set to the default 0.8, which is recommended for optimal analysis sensitivity and stringency.

Enrichment analyses were run using two different methods. First, an exploratory assessment, IPA Core Expression analysis was run using IPA microRNA target filter output. The IPA microRNA target filter queries the TargetScan, TarBase, and miRecords databases, which contain predicted and experimentally validated miRNA-mRNA interactions at all prediction confidence levels. Enriched Canonical Pathways and Diseases/Biological Functions were identified using an uncorrected significance threshold of 0.05. Full IPA analysis settings are available in supplementary materials (File S1). Second, a more stringent approach, the DIANA miRPath tool, v.3 (Vlachos et al., 2015) was used to identify enriched KEGG pathways from transcripts identified by the microT-CDS algorithm. miRPath was run using the “Pathways Union” algorithm, with “Unbiased Empirical Distribution” selected as the enrichment method. In contrast to the commonly used miRPath “Genes Union” setting, which is similar to the approach taken by the IPA Core analysis, Pathways Union utilizes an FDR corrected, merged p-value that represents the likelihood a certain biological pathway is targeted by at least one of the input miRNAs. Further, use of an empirical distribution, which includes all transcripts predicted to be targets of miRNAs (vs. the entire murine transcriptome), has been suggested to reduce algorithm bias and increase prediction accuracy (Bleazard, Lamb, & Griffiths-Jones, 2015).

### 3.5 Statistical Analysis

Behavioral data were analyzed in SPSS (IBM, Armonk, NY) with factorial analysis of variance (ANOVA). Independent factors included stress (Control vs CVSS), strain (C57BL/6J vs BALB/cJ), sex (male vs female), day (1-11), concentration (morphine: 0.3/0.7/1 or quinine: 0.2/0.4/0.55 mg/ml), and drug group (chronic saline vs chronic drug) where appropriate. Interactions that included multiple variables were interpreted with successive ANOVAs with fewer factors. Tukey’s *post hoc* test was used and the alpha level set at 0.05.

## 4 Results

### 4.1 Morphine Sensitization

The impact of adolescent social stress on morphine sensitization varied by strain. Adult C57BL/6J mice exposed to adolescent social stress had blunted morphine sensitization, but BALB/cJ mice did not (Fig 1B,D vs. 1F,H). In the overall analysis with strain, stress, sex, drug group, and day as factors, we observed many significant effects involving strain ranging from a main effect of strain to 4-way interactions involving this factor, thus we conducted further analyses split by strain (see Supplementary Table 2 for full ANOVA results).

**Figure 1:**
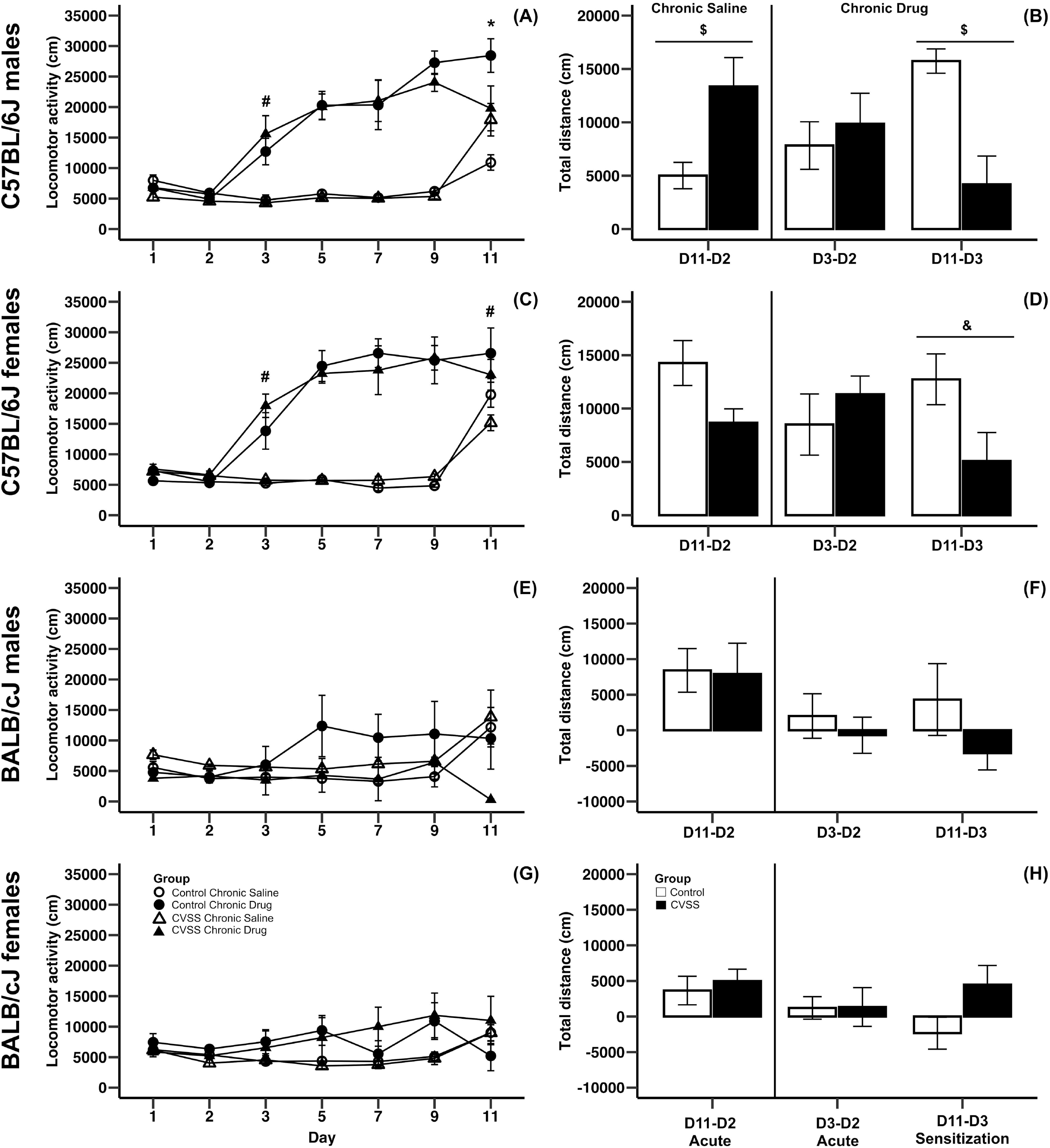
Morphine-induced acute behavioral stimulation and sensitization in adult C57BL/6J and BALB/cJ mice. Daily mean locomotor distance (cm) ± SEM are presented in panels A,C,E,G: (A) C57BL/6J males, (C) C57BL/6J females, (E) BALB/cJ males, and (G) BALB/cJ females. Within-subject initial stimulant response to morphine (mean ± SEM) are presented in panels B,D,F,H: (B) C57BL/6J males, (D) C57BL/6J females, (F) BALB/cJ males, and (H) BALB/cJ females. Chronic saline animals are on the left side and chronic drug treated mice on the right side in each panel, with sensitization in the chronic drug treated mice shown in the most right-hand bars. N = 5-8 per stress and drug condition. *p<0.05 for stress X drug group interaction; #p<0.05 for the main effect of drug group; $p<0.05 for the main effect of stress; &p=0.055 for the main effect of stress

In C57BL/6J mice, we observed multiple significant effects (see Table 3 for full results) including a day X stress X sex X drug group interaction (F_6,264_ = 2.5, p < 0.05). To decompose this 4-way interaction, follow up analyses were conducted in each sex separately. In male C57BL/6J mice, we observed a significant main effect of day (F_6,132_ = 38.4, p < 0.001), drug group (F_1,22_ = 88.2, p < 0.001), day X drug group (F_6,132_ = 22.5, p < 0.001), and day X stress X drug group interactions (F_6,132_ = 4.3, p < 0.01). The interactions between day and group are consistent with the experimental design and suggest that the pattern of activity across days is dependent on the experimental drug group. Thus, further analyses were limited to the key days that examined between-groups acute locomotor stimulation (Day 3) and sensitization (Day 11) (Gubner et al., 2014; Miller and Kamens, 2019). On Day 3, the first day of morphine exposure in the chronic drug group, C57BL/6J mice showed increased locomotion in response to morphine injection compared to the chronic saline group that received saline, indicating the expected morphine stimulant response (main effect of drug group: F_1,22_ = 24.4, p < 0.01). There were no other main effects or interactions with stress on Day 3. On Day 11, the 5^th^ day of morphine treatment in the chronic drug group compared to the first day of morphine exposure in the chronic saline group, there was a significant main effect of drug group (F_1,22_ = 11.7, p < 0.01) and a stress X drug group interaction (F_1,22_ = 7.6, p < 0.05). *Post hoc* analyses demonstrated that for control animals there was a significant difference between the chronic drug and chronic saline groups (p < 0.05), indicating the expected morphine sensitization. In contrast, there was no significant difference between the chronic drug and chronic saline groups in the CVSS animals, suggesting attenuated morphine sensitization.

**Table 2.** Treatment protocol for morphine sensitization protocol.

| Group | Day 1 | Day 2 | Day 3 | Day 4 | Day 5 | Day 6 | Day 7 | Day 8 | Day 9 | Day 10 | Day 11 |
| --- | --- | --- | --- | --- | --- | --- | --- | --- | --- | --- | --- |
| *Chronic Saline* |  |  |  |  |  |  |  |  |  |  |  |
| Injection | Saline | Saline | Saline | None | Saline | None | Saline | None | Saline | None | MO |
| Test | Yes | Yes | Yes | No | Yes | No | Yes | No | Yes | No | Yes |
| *Chronic Drug* |  |  |  |  |  |  |  |  |  |  |  |
| Injection | Saline | Saline | MO | None | MO | None | MO | None | MO | None | MO |
| Test | Yes | Yes | Yes | No | Yes | No | Yes | No | Yes | No | Yes |

MO: Morphine.

**Table 3.**
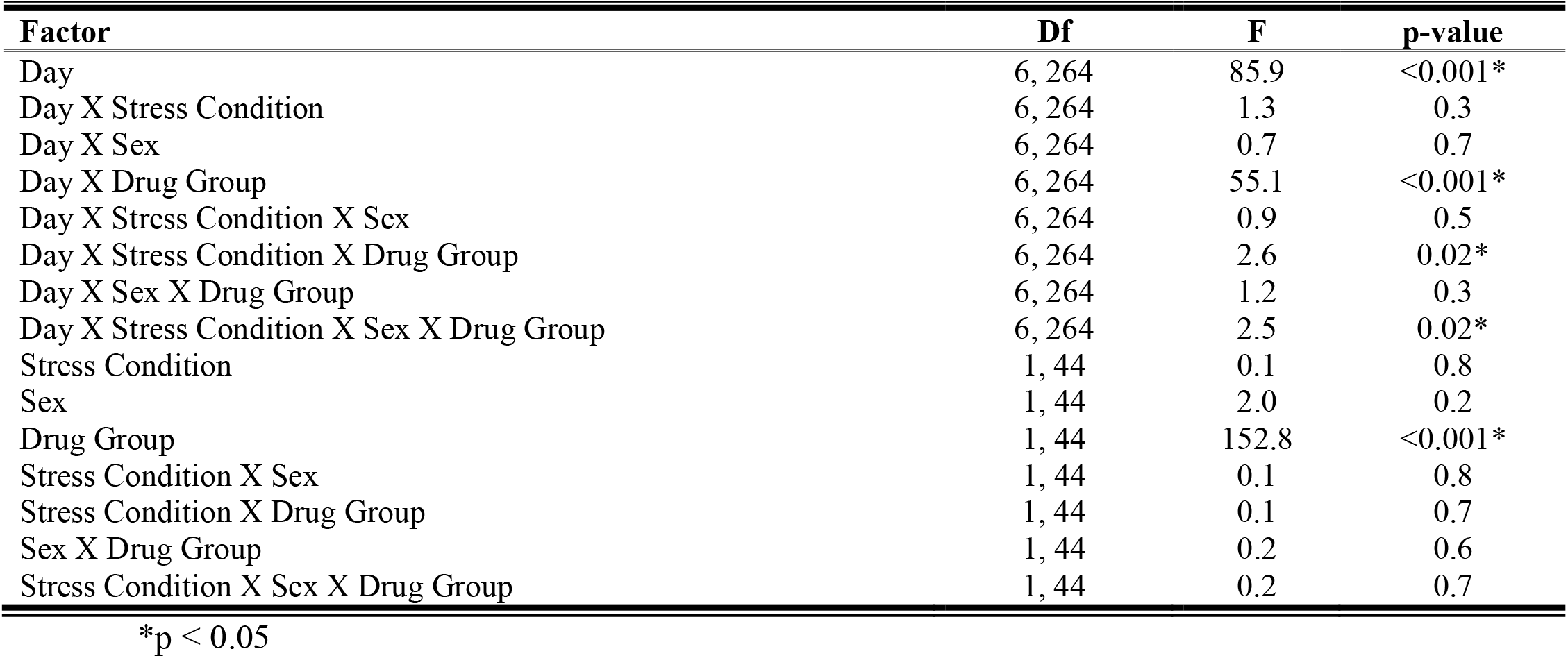
Morphine Sensitization ANOVA Output from C57BL/6J mice including Day, Stress Condition, Sex, and Drug Group as independent factors.

A similar pattern was observed in male C57BL/6J mice with the within-group analyses. In chronic saline exposed male C57BL/6J mice, CVSS males had a more robust acute locomotor response to morphine compared to unstressed control males (Day 11 – Day 2; t_11_ = -2.6, p < 0.05; Fig 1B). But, there was no significant difference in the acute locomotor response to morphine between stress and control animals in the chronic drug groups (Day 3 – Day 2). Findings from the within-group measure of sensitization (Day 11 – Day 3), replicated the between-group analysis (Day 11 above). Control male C57BL/6J mice exposed to chronic morphine exhibited robust behavioral sensitization that was attenuated in CVSS animals (t_11_ = 3.8, p < 0.01; Fig 1B).

In female C57BL/6J mice, we observed a significant main effect of day (F_6,132_ = 48.7, p < 0.001), group (F_1,22_ = 68.4, p < 0.001), and a day X drug group interaction (F_6,132_ = 34.3, p < 0.001). Importantly, in female C57BL/6J mice the main effect of stress and interactions with this variable were not significant. When examining days indicative of acute stimulation (Day 3) and sensitization (Day 11) there were significant main effects of drug group (t_24_ = -5.5, p < 0.001, t_24_ = -2.4, p < 0.05, respectively). Compared to chronic saline mice, chronic drug mice exhibited greater locomotor response on Day 3 (mean ± SEM, 5541 ± 258, 15732 ± 1863, respectively), indicative of the acute stimulant response, and a greater response on Day 11, indicative of between-groups sensitization (mean ± SEM, 17994 ± 1463, 24929 ± 2475, respectively). In the within-group analysis, no significant stress group differences were observed for the acute locomotor response in either the chronic saline or chronic drug groups. When behavioral sensitization was examined, there was a statistical trend (p = 0.055) where female C57BL/6J mice exposed to chronic adolescent social stress had blunted morphine sensitization compared to control animals (Fig 1D).

In BALB/cJ mice, the 100 mg/kg dose of morphine did not result in locomotor stimulation or sensitization (Fig 1E-H). The overall analysis within this strain demonstrated a main effect of day (F_6,252_ = 4.9, p < 0.001) and a day X drug group interaction (F_6,252_ = 6.6, p < 0.001). *Post hoc* analysis found no significant effects on Day 3 or Day 11. Further, there were no significant effects observed in the within-group analysis for either acute stimulation or sensitization in BALB/cJ mice.

### 4.2 Morphine Consumption

Five C57BL/6J mice (spread among stress conditions and sex) died during the consumption study. Veterinarian examination revealed that the animals did not have food in their stomachs suggesting drug-induced anorexia. Although we followed a published 24-hour morphine access consumption paradigm (Eastwood and Phillips, 2014), future work should consider limiting drug availability to 18 h to decrease the chance of such an adverse effect (Kamens et al., 2005).

Adolescent social stress did not influence adult morphine consumption in either C57BL/6J or BALB/cJ (Fig 2) mice. In an overall analysis of morphine consumption (mg/kg) there was a significant main effect of strain (F_1,95_ = 180.9, p < 0.001) and a strain X concentration interaction (F_2,190_ = 43.5, p < 0.001), thus in further analyses C57BL/6J and BALB/cJ mice were analyzed independently. In C57BL/6J mice, morphine consumption increased in a concentration dependent manner (Fig 2A; main effect of concentration: F_2,90_ = 348.8, p < 0.001, all *post hoc* p < 0.001), but no other significant main effects or interactions were observed. There were no significant effects on quinine consumption (Fig 2C). For total fluid intake, while there was a significant sex X concentration interaction (F_2,90_ = 3.3, p < 0.05), *post hoc* comparison suggest this was driven by a trend (p = 0.06) for the males to consume more fluid compared to females (8.9 ± 0.6, 7.7 ± 0.3, respectively), but only when the middle morphine concentration (0.7 mg/ml) was available.

**Figure 2:**
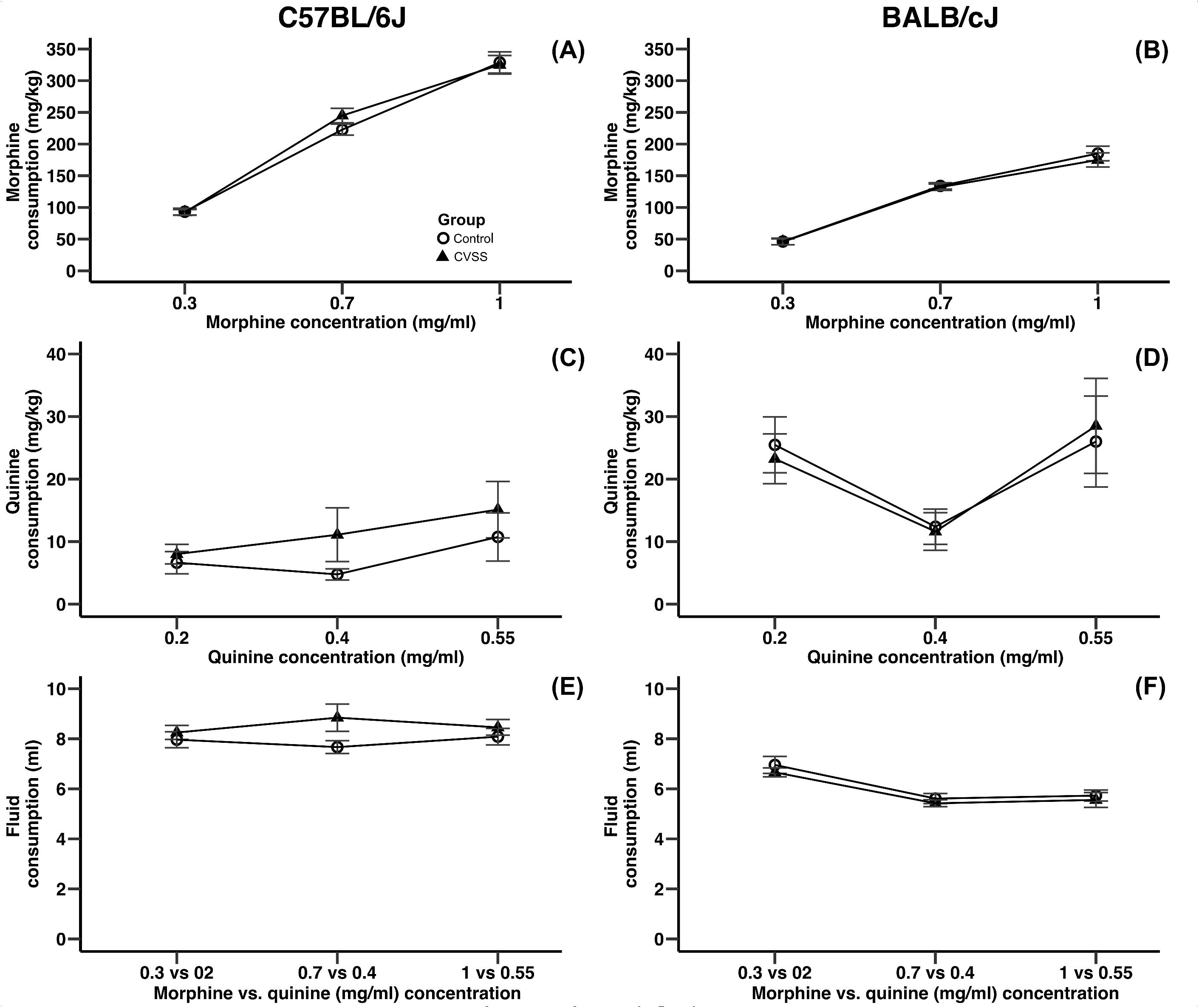
Adolescent stress did not influence morphine or quinine consumption in adult C57BL/6J or BALB/cJ mice. Morphine consumption and quinine consumption (mean±SEM) in (A,C) C57BL/6J and (B,D) BALB/cJ mice. Mean (±SEM) fluid intake in (E) C57BL/6J and (F) BALB/cJ mice. N =22–28 per strain and stress condition.

Similar to the results observed in C57BL/6J mice, in BALB/cJ mice drug concentration influenced intake, but no main effects or interactions with stress condition were observed. BALB/cJ mice consumed significantly more morphine when the 0.7 and 1 mg/ml morphine concentrations were available compared to the 0.3 mg/ml concentration (Fig 2B, main effect of concentration: F_2,100_ = 228.4, p < 0.001, all *post hoc* p < 0.001). Importantly, there were no main effects or interactions with stress condition. In BALB/cJ mice, there was a significant main effect of concentration on quinine consumption (F_2,100_ = 6.0, p < 0.01). Here, quinine consumption was lower when the middle quinine concentration was available relative to the low or high concentrations (Fig 2D, all *post hoc* p < 0.05). Finally, when total fluid intake was examined there was a significant main effect of concentration (F_2,100_ = 27.2, p < 0.001) and sex (F_1,50_ = 4.9, p < 0.05). BALB/cJ mice drank significantly more fluid when the 0.3 mg/ml (6.8 ± 0.2) morphine concentration was available relative to both the 0.7 (5.5 ± 0.1) or 1 (5.6 ± 0.2) mg/ml concentrations (Fig 2F, all *post hoc* p < 0.001). Overall, male BALB/cJ mice consumed more fluid than female BALB/cJ mice (6.2 ± 0.1, 5.7 ± 0.2, respectively).

### 4.3 Acute Stress

Adult C57BL/6J mice exposed to adolescent social stress had a slower recovery of CORT levels following an acute restraint stress compared to control mice (Fig. 3). Baseline levels of CORT were not influenced by stress or sex (data not shown). We then examined stress reactivity (30 min CORT – baseline CORT) and again no significant effects were observed. In contrast, when recovery (90 min CORT – 30 min CORT) was examined there was a significant main effect of stress (F_1,24_=12.1, p<0.01) and sex (F_1,24_=4.9, p<0.05), but the interaction of these variables was not significant. Mice exposed to CVSS had a slower CORT recovery compared to control animals (3.3 ± 32.5, -88.7 ± 32.2, respectively) and females recovered more slowly than males (45.1 ± 26.9, -99.5 ± 28.5, respectively).

**Figure 3:**
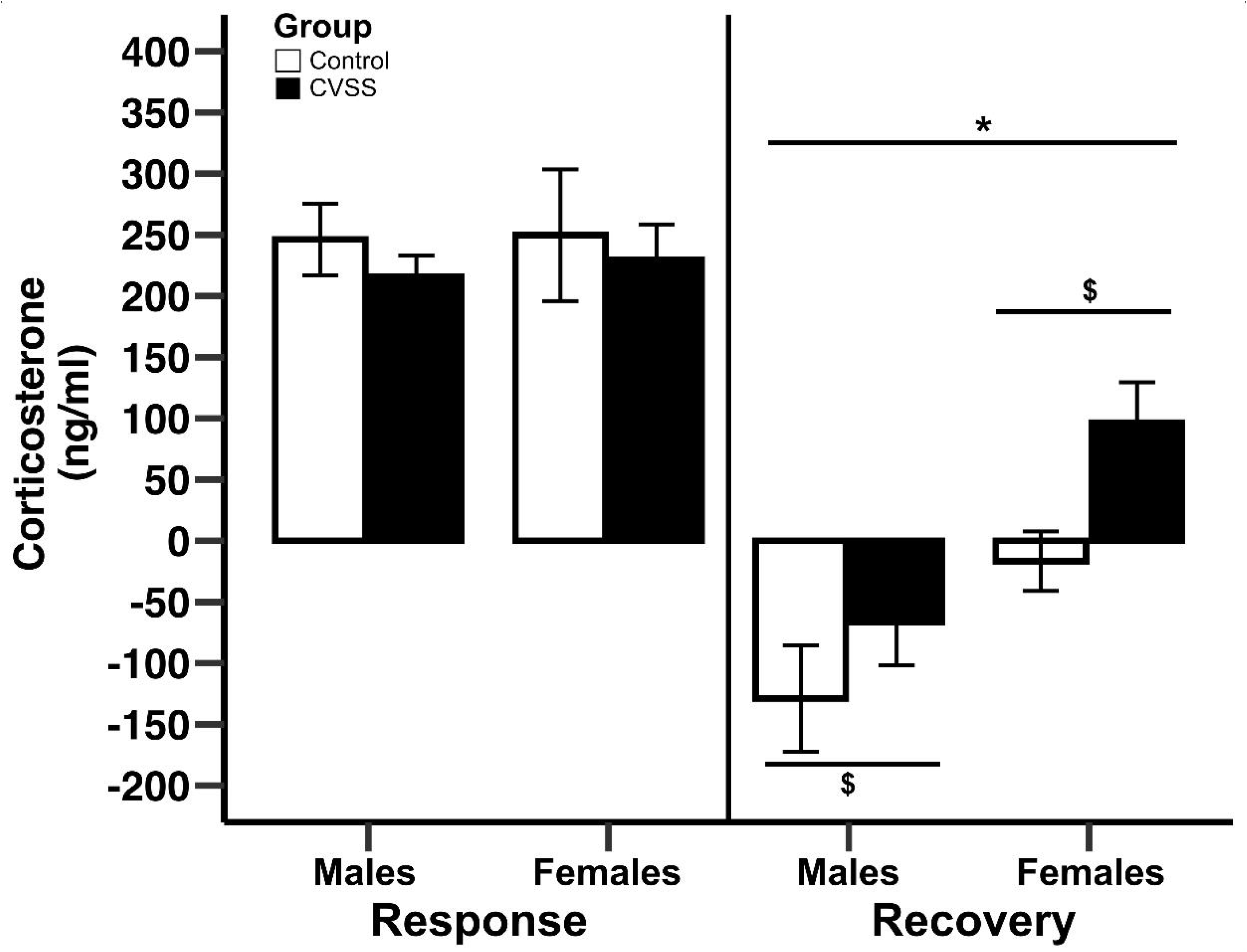
Change in corticosterone following 15 minutes of restraint stress in adult C57BL/6J mice. Mean corticosterone (ng/ml) ± SEM in C57BL/6J male and female mice N = 14 per stress condition. *p<0.05 for main effect of sex; $p<0.05 for main effect of stress group

### 3.4 Prefrontal Cortex (PFC) miRNA Gene Expression

In adult C57BL/6J male mice of the 904 miRNA identified through RNA sequencing, twelve (miR-429-3p, miR-200a-3p, miR-96-5p, miR-141-3p, miR-200b-3p, miR-183-5p, miR-200a-5p, miR-182-5p, miR-200c-3p, miR-141-5p, miR-183-3p, miR497b) were downregulated following adolescent social stress (for full results see supplementary File S2). The DIANA microT-CDS algorithm, run with the 12 differentially expressed miRNA, predicted 5433 total miRNA-mRNA relationships, with 2820 unique mRNA targets. Five significant KEGG pathways were identified by miRPath: MAPK signaling, Glycosphingolipid biosynthesis (lacto and neolacto series), AMPK signaling, Glycosphingolipid biosynthesis (globo series), and Gap junction (see also Table 4). At a significance threshold of 0.05, IPA Core Expression analysis identified 500 enriched Diseases/Biological Functions and 267 enriched Canonical Pathways. See Table 5 for the top five enrichment terms in each category and supplementary Files S3-S4 for full IPA output.

**Table 4.** miRPath output.

| KEGG pathway | p-value | #genes | #miRNAs |
| --- | --- | --- | --- |
| MAPK signaling pathway | 0.015 | 5 | 1 |
| Glycosphingolipid biosynthesis – lacto and neolacto series | 0.018 | 2 | 1 |
| AMPK signaling pathway | 0.027 | 4 | 1 |
| Glycosphingolipid biosynthesis – globo series | 0.027 | 3 | 2 |
| Gap junction | 0.048 | 2 | 1 |

**Table 5.** IPA output.

| IPA top 5 canonical pathways |  | p-value |
| --- | --- | --- |
| Molecular mechanisms of cancer |  | 3.98E-11 |
| BMP signaling pathway |  | 1E-10 |
| Synaptogenesis signaling pathway |  | 2.75E-10 |
| Cardiac hypertrophy signaling |  | 3.24E-09 |
| GNRH signaling |  | 1.66E-08 |

| IPA disease/function annotation | p-value | #molecules |
| --- | --- | --- |
| Non-hematological solid tumor | 7.44E-208 | 4133 |
| Nonhematologic malignant neoplasm | 1.25E-204 | 4118 |
| Cancer | 3.74E-190 | 4164 |
| Malignant solid tumor | 1.4E-189 | 4151 |
| Non-melanoma solid tumor | 5.4E-189 | 4079 |

## 5 Discussion

In the present study we examined the effect of adolescent social stress on adult morphine behaviors in two inbred strains of mice, and characterized two key biological mechanisms that may account for decreased morphine sensitization in C57BL/6J mice exposed to adolescent social stress. We found that adolescent social stress blunted morphine sensitization in adult C57BL/6J, but not BALB/cJ, mice, an effect that was more robust in male mice. This behavioral effect of stress emerged after repeated drug administration, as there was little evidence of a change in the acute locomotor response to morphine. Adolescent stress did not influence adult voluntary morphine consumption in either strain. We observed long-term changes in both the recovery of the CORT response following an acute stressor and in the expression of adult miRNA in the prefrontal cortex. It is possible that these changes reflect mechanisms linked to the altered morphine response. These data indicate that a gene-environment interaction may influence opioid behaviors.

Our results indicate that male C57BL/6J mice are particularly vulnerable to lasting effects of adolescent stress. In both the within- and between-groups analysis we observed that control animals developed normal morphine-induced behavioral sensitization, but this was blunted in animals that were exposed to social stress in adolescence. A similar statistical trend (p = 0.055) was observed in female C57BL/6J mice with the within-group analysis. Due to the complex design of this experiment, there were 5-8 mice per sex, stress condition, and drug group; thus, it is possible we were underpowered to detect significant effects. The fact that there is a known statistical power advantage of within-subject design may explain why we observed a trend in the female C57BL/6J mice using this design (Keppel, 1991).

Our findings that adolescent stress blunts morphine sensitization aligns with two other preclinical studies. In work by Courdereau, no morphine conditioned place preference (8, 16, 64, or 100 mg/kg) was observed in male NMRI mice that were exposed to 30 days of social isolation that started in adolescence (Coudereau et al., 1997). Similar abolishment of morphine conditioned place preference was observed in male Lister hooded rats that were isolated for 6 weeks starting at weaning (Wongwitdecha and Marsden, 1996). Both of these studies incorporated isolation stress similar to ours, but it is possible that these effects extent to other stressors as well.One limitation of the current work is that we utilized only a single high dose of morphine, thus we can’t determine if adolescent stress shifts sensitivity to morphine. A similar series of experiments with additional morphine doses may provide some insight, but when our results are compared to earlier findings with morphine conditioned place preference (Wongwitdecha and Marsden, 1996; Coudereau et al., 1997) it could be hypothesized that adolescent stress blocks normal behavioral responses to opioids. Obviously, additional work is necessary to confirm this hypothesis.

The differences in morphine behavioral responses appear to emerge after repeated drug exposures as we saw little indication that stress altered the acute locomotor response to morphine. We observed that CVSS increased morphine stimulation in the male C57BL/6J chronic saline group, but not in the chronic drug group or with the between-groups analysis. These findings are consistent with a previous study that demonstrated that 14 days of restraint stress in adult C57BL/6J mice had no effect on acute morphine locomotor stimulation (Xu et al., 2014). However, these results differ from prior research in adult rats that have demonstrated that stress (restraint, handling, social defeat, foot shock, or social isolation) potentiates morphine-induced locomotor stimulation (Leyton and Stewart, 1990; Deroche et al., 1994; Stöhr et al., 1999). The stressors in these studies varied in duration and number of exposures. Stöhr and colleagues demonstrated that one 60-min restraint stress was not enough to potentiate morphine locomotor stimulation, but 3 exposures to restraint stress, handling or social defeat did. The stressors in the other papers were longer with 5 sessions of 20 min foot shock or 17 days of continuous social isolation in adulthood. These procedural differences make it difficult to draw strong conclusions, but it is interesting to note that while significant effects of stress exposure on morphine stimulation have been observed in rats, there is currently little evidence in C57BL/6J mice.

In the present work, we observed no difference in morphine consumption following adolescent social stress. Our findings align with work in adult male C57BL/6J mice. Exposure to chronic social stress increased morphine preference immediately (1 day) after stress exposure, but the effect was no longer present when tested 14 days post-stress exposure (Cooper et al., 2017). In the current experiment, mice were given access to morphine beginning 8-11 days following the stress exposure. We originally hypothesized that stress during adolescence would result in long-lasting differences in drug consumption because of the critical time in brain development, but that was not observed with morphine consumption. Instead the effects of stress on opioid consumption may be time-dependent and only evident immediately after stress exposure. This is important when considering factors that could lead to drug use initiation.

In both the morphine sensitization and consumption experiments, we found pronounced differences between C57BL/6J and BALB/cJ mice. These differences align with prior research demonstrating strain differences in morphine behaviors (Belknap et al., 1993; Semenova et al., 1995; Kest et al., 2002; Kennedy et al., 2011). For example, consistent with the prior literature C57BL/6J mice consumed significantly more morphine than BALB/cJ mice (Belknap et al., 1993). In the current study our paradigm produced robust behavioral sensitization in C57BL/6J mice, but not in BALB/cJ mice. It is possible that the dose of morphine (100 mg/kg) we chose was too high for this strain. While we found no effects of stress in BALB/cJ mice, it is critical to recognize that different effects may be observed on other morphine behaviors or in response to other doses of morphine used in our paradigms.

Here we show that exposure to chronic social stress in adolescence changes the physiological response to an acute stressor in adulthood. In particular, we found that it took stressed mice longer to recover following a 15 min restraint stress. This finding is in line with research suggesting that animals chronically exposed to one type of stressor beforehand have an altered hormonal response to a novel stressor (Romeo, 2013). For example, adolescent rats exposed to 7 days of a cold room stressor then restraint stress had a delayed recovery of CORT to the restraint stress. In contrast, this effect was not observed in adult rats or in adolescent rats exposed to 8 days of restrain stress (Lui et al., 2012). In this prior study, all stress exposures occurred during adolescence, but the current findings suggest that this altered physiological response to an acute stressor may last into adulthood.

In this study we observed 12 miRNA in the adult prefrontal cortex that were downregulated following adolescent social stress. We chose to examine differential gene expression in the prefrontal cortex because of our prior work which demonstrated lasting changes in neuronal excitability in this region (Caruso et al., 2018a). A differentially expressed miRNA of interest that we identified was miR-183. This finding is consistent with a prior study that showed chronic immobilization stress in adulthood decreased the expression of this gene in the amygdala and CA1 region of the hippocampus (Meerson et al., 2010). Interestingly, in that study an upregulation was identified after acute immobilization stress, suggesting that the duration of stress exposure is important. Time-dependent effects of stress exposure on other miRNAs have also been observed. For example, miR-200a-3p was shown to be down regulated in the prefrontal cortex of rats exposed to 4 weeks of chronic unpredictable stress in adulthood; a finding similar to our own. In contrast, after 12 weeks of stress exposure this same miRNA was upregulated (Satyanarayanan et al., 2019). We believe that this investigation produced the first known data to examine the global patterns of prefrontal cortex miRNA expression following adolescent stress. Given that these small RNA molecules regulate mRNA expression, our findings could contribute to the altered behavioral responses we observed, but additional work is necessary to confirm this hypothesis. Further, these data provide additional support that repeated exposure to stress in adolescence may compromise typical development of the PFC.

Functional enrichment analysis of the differentially expressed miRNA and their predicted mRNA targets identified proCancer-related pathways among the top enrichment terms in both miRPath and IPA analyses (Supplementary Table 4 & 5). Cancer-related signaling tends to involve ubiquitous molecules and pathways that often also participate in neurological functioning. For example, the top enriched miRPath KEGG pathway was MAPK signaling (Table 4), a biological pathway well known to mediate cancer formation as well as learning and memory (Atkins et al., 1998; Wagner and Nebreda, 2009). Further, in addition to their known role in tumor formation (Inokuchi et al., 1987), glycosphingolipids have recently been implicated in addiction (Kalinichenko et al., 2018). IPA core analysis identified overall top enrichment terms related to cancer and synaptogenesis (Supplementary Table 5). It is also notable that the top IPA Diseases/Functions terms under the subcategory “Physiological System Development and Function” was “Nervous System Development and Function.” Thus, adolescent stress appears to alter expression of adult miRNAs broadly regulating signaling pathways related to cancer and nervous system development/functioning. These specific miRNA and pathways serve as mechanisms by which adolescent stress my change later opioid behaviors.

The current work demonstrates a gene-environmental interaction that predisposes certain individuals to opioid behaviors following adolescent stress. Specifically, adult C57BL/6J mice exposed to adolescent social stress had attenuated morphine sensitization compared to control animals, whereas BALB/cJ mice did not. It remains possible adolescent stress may impact morphine sensitization in BALB/cJ if a more optimal dose of this drug was used in this strain. This is not the first time we have observed such a strain X stress dependent effects with this stressor; a similar finding was seen with nicotine behaviors, with altered sensitivity to nicotine in male BALB/cJ, but not C57BL/6J mice (Caruso et al., 2018b, 2018a). Given that the majority of addiction research utilizes C57BL/6J mice it is important to recognize that we may miss gene-environment interactions if utilizing only a single mouse strain.

## Conflict of Interest

The authors declare that the research was conducted in the absence of any commercial or financial relationships that could be construed as a potential conflict of interest.

## Ethics Statement

This study was approved (protocol 00246) and carried out in accordance with the recommendations of the Penn State University Institutional Care and Use Committee.

## Author Contributions

HK, CM, JC, and SC designed the study. HK, CM, and JC performed the experiments. HK, CM, JC, DZ, WH, CS, AS, IA, and SC analyzed the data. HK, WH, TG, DF, PG, and SAC provided conceptual input into the project and edited the manuscript. HK, JC, and DZ did the majority of the writing.

## Funding

This work was supported by the National Institutes of Health grants P50 DA039838 and UL1 TR002014. Additional support came from The Pennsylvania State University Biobehavioral Health Department, the Consortium to Combat Substance Abuse, and the Social Science Research Institute. The funders had no role in study design, data collection and analysis, decision to publish, or preparation of the manuscript. The content is solely the responsibility of the authors and does not necessarily represent the official views of the funding institutions as mentioned above.

## Acknowledgments

The authors would like to thank Rachel Bartlett, Sophia Kenney, Aidan Peat, Lan-Nhi Phung, Colton Ruggery, Cassandra Secunda, Kylie Stuhltrager, and Grant Swisher for help with this project.

## Data Availability Statement

The data that support the findings of this study are available from the corresponding author upon reasonable request. Sequencing data are available from the NCBI GEO database (experimental series accession number: to become available upon publication acceptance).

## Notes

### Competing Interest Statement

The authors have declared no competing interest.

