## Supplemental Table 2 for "Effect of Adolescent Stress on Adult Morphine-Induced Behavioral Sensitization is Dependent Upon Genetic Background"

**Supplemental Table 3.** Full Morphine Sensitization ANOVA Output including Day, Strain, Stress Condition, Sex, and Drug Group as Independent Factors.

| **Factor** | **df** | **F** | **p-value** |
| --- | --- | --- | --- |
| Day | 6, 516 | 58.5 | <0.001* |
| Day X Strain | 6, 516 | 22.4 | <0.001* |
| Day X Stress Condition | 6, 516 | 0.9 | 0.4 |
| Day X Sex | 6, 516 | 0.1 | 1 |
| Day X Drug Group | 6, 516 | 44.1 | <0.001* |
| Day X Strain X Stress Condition | 6, 516 | 0.6 | 0.7 |
| Day X Strain X Sex | 6, 516 | 0.8 | 0.6 |
| Day X Strain X Drug Group | 6, 516 | 14.6 | <0.001* |
| Day X Stress Condition X Sex | 6, 516 | 0.3 | 0.9 |
| Day X Stress Condition X Drug Group | 6, 516 | 1.4 | 0.2 |
| Day X Sex X Drug Group | 6, 516 | 0.2 | 1.0 |
| Day X Strain X Stress Condition X Sex | 6, 516 | 2.4 | 0.03* |
| Day X Strain X Stress Condition X Drug Group | 6, 516 | 1.1 | 0.4 |
| Day X Strain X Sex X Drug Group | 6, 516 | 1.5 | 0.2 |
| Day X Stress Condition X Sex X Drug Group | 6, 516 | 3.0 | 0.006* |
| Day X Strain X Stress Condition X Sex X Drug Group | 6, 516 | 1.0 | 0.4 |
| Strain | 1, 86 | 72.7 | <0.001* |
| Stress Condition | 1, 86 | 0.3 | 0.6 |
| Sex | 1, 86 | 1.4 | 0.2 |
| Drug Group | 1, 86 | 69.9 | <0.001* |
| Strain X Stress Condition | 1, 86 | 0.04 | 0.8 |
| Strain X Sex | 1, 86 | 0.3 | 0.6 |
| Strain X Drug Group | 1, 86 | 43.6 | <0.001* |
| Stress Condition X Sex | 1, 86 | 0.5 | 0.5 |
| Stress Condition X Drug Group | 1, 86 | 1.4 | 0.2 |
| Sex X Drug Group | 1, 86 | 1.7 | 0.2 |
| Strain X Stress Condition X Sex | 1, 86 | 0.2 | 0.7 |
| Strain X Stress Condition X Drug Group | 1, 86 | 0.6 | 0.4 |
| Strain X Sex X Drug Group | 1, 86 | 0.5 | 0.5 |
| Stress Condition X Sex X Drug Group | 1, 86 | 2.9 | 0.09 |
| Strain X Stress Condition X Sex X Drug Group | 1, 86 | 1.5 | 0.2 |

*p < 0.05
